# Minimally Complex Nucleic Acid Feedback Control Systems for First Experimental Implementations

**DOI:** 10.1101/867945

**Authors:** Nuno M. G. Paulino, Mathias Foo, Tom F. A. de Greef, Jongmin Kim, Declan G. Bates

**Affiliations:** Warwick Integrative Synthetic Biology Centre, School of Engineering, University of Warwick, Coventry CV4 7AL, UK (,); School of Mechanical, Aerospace and Automotive Engineering, Coventry University, Coventry CV1 5FB, UK; Department of Biomedical Engineering, Eindhoven University of Technology (TUE), Eindhoven, The Netherlands; Department of Integrative Biosciences and Biotechnology, Pohang University of Science and Technology (POSTECH), Pohang, Gyeongbuk, 37673, South Korea

**Keywords:** Feedback control, Chemical reaction networks, Nucleic acids, Strand Displacement Circuits, Synthetic Biology

## Abstract

Chemical reaction networks based on catalysis, degradation, and annihilation may be used as building blocks to construct a variety of dynamical and feedback control systems in Synthetic Biology. DNA strand-displacement, which is based on DNA hybridisation programmed using Watson-Crick base pairing, is an effective primitive to implement such reactions experimentally. However, experimental construction, validation and scale-up of nucleic acid control systems is still significantly lagging theoretical developments, due to several technical challenges, such as leakage, crosstalk, and toehold sequence design. To help the progress towards experimental implementation, we provide here designs representing two fundamental classes of reference tracking control circuits (integral and state-feedback control), for which the complexity of the chemical reactions required for implementation has been minimised. The supplied ‘minimally complex’ control circuits should be ideal candidates for first experimental validations of nucleic acid controllers.

## 1. INTRODUCTION

Synthetic circuits implemented with DNA strand displacement (DSD) reactions are biologically compatible, programmable, and have been demonstrated *in vivo* (Hemphill and Deiters (2013)), making DSD networks promising candidates for the construction of synthetic feedback controllers in biomolecular environments. Feedback control systems can be represented with elementary chemical reaction networks (CRNs), which provide an abstract layer for the design of mathematical operators (Buisman et al. (2008)), before being translated to biomolecular applications. Networks of catalysis, degradation and annihilation reactions can then be mapped into equivalent DSD reactions (Soloveichik et al. (2010)), where the sequences of nucleotides in the DNA strands effectively program the biochemical interactions and the rates in each strand displacement reaction (Chen et al. (2013), Zhang et al. (2018)).

Concentrations of biomolecular species are at first glance ill-suited to represent signals in feedback control theory, since they are restricted to positive quantities. An approach that circumvents this limitation is the so-called dual rail representation of system state variables with pairs of concentrations, as in Seelig et al. (2006), where a signal *x*(*t*) is represented by the difference between two concentrations *x*(*t*) = *x*^+^(*t*) − *x*^−^(*t*), corresponding to two chemical species *X*^+^ and *X*^−^. This dual rail framework provides a representation of negative signals, which are necessary for computing the error in a negative feedback control system, and enables the use of molecular concentrations to design feedback control circuits based on linear operators and frequency or state-space designs, see examples in Oishi and Klavins (2011); Yordanov et al. (2014); Chiu et al. (2015); Foo et al. (2017); Paulino et al. (2019).

Examples of successful experimental implementations of nucleic-acid feedback control circuits have yet to emerge, however. DSD networks implementing open-loop cascades for logical or analogue computation have been reported (Thubagere et al. (2017)), but there is a significant theory-experiment gap for dynamical circuits implementing negative feedback, where the transient dynamics impact system stability and performance, for a variety of reasons.

Although it is possible to predict the toehold affinities from their nucleotide sequence (Zhang et al. (2018)), the kinetics are altered by unwanted bindings, which modify the reaction rate constants and the dynamics (see, for example, the oscillating circuits in Srinivas et al. (2017) or the seesaw gate reported in Qian and Winfree (2011)).

The triggering of undesired reactions is another key experimental challenge, which leads to leakage of outputs in the absence of input. Potential methods of mitigation include *clamps* which impede the spurious hybridisations (as in Srinivas et al. (2017) and Wang et al. (2017)), or compartmentalisation and localisation strategies as proposed in Dannenberg et al. (2015), Chatterjee et al. (2017), and Joesaar et al. (2019), which keep apart strand complexes that may trigger leak reactions. Furthermore, leakage is aggravated at high concentrations, and therefore the reacting species are typically kept at low concentrations (Wang et al. (2018)), which together with limits on hybridisation rates (Zhang et al. (2018)), places upper bounds on the speed of these circuits. Localisation can also help here, by placing adjacent gates, which can interact without diffusion at faster rates, as discussed in Qian and Winfree (2011). Finally, even if spurious reactions are avoided, it may still be necessary to manage the sequence of reactions, which compete for common reactants, either through compartmentalisation, or with timescale separation (as adopted in Cherry and Qian (2018)).

All of the above experimental challenges can be mitigated by designs that reduce the number of reactions in the circuit. Fewer reactions decrease the number of designs for the template strands, which demands less work to characterise and tune the kinetics. Fewer species also decreases the chance of unwanted interactions and leakage. The issue is particularly relevant in feedback systems employing the dual rail representation of negative signals, which requires a duplication of most of the reactions (as discussed in Foo et al. (2017)).

A key challenge for theorists is therefore to design the least complex circuits, with the fewest number of chemical reactions, which still accurately represent a given negative feedback control problem, in order to maximise the likelihood of successful experimental implementation with currently available technical capabilities. To this end, minimally complex representations of two fundamental classes of feedback control systems are proposed here. The first applies integral control for reference tracking, with a single tuning parameter, to a stable first order system. The second example accomplishes reference tracking and stabilisation of the classic double integrator plant through static state feedback of the two integrated states. The controller has two design parameters, and the open-loop system is marginally stable with two poles at the origin. Both circuits operate within the dual rail framework, and parameterise control requirements like steady state tracking and transient dynamics.

Note that, by representing the gains and subtractions with individual CRNs, the state feedback scheme in Paulino et al. (2019) is already simplified with respect to a classical PID controller, since it combines the gains with the subtraction in the same CRN. Here we reduce the complexity further by combining gain, integration and subtraction in the same chemical reactions. This allows us to remove two catalysis, two degradation, and one annihilation reactions in the integral control problem, with respect to the approach in Paulino et al. (2019). In the state feedback controller, we remove four catalysis and one annihilation reactions, by combining the subtraction with the integration of the first state, and by selecting a plant that does not need degradation reactions for its representation. As we will show, both problems may be implemented using similar low numbers of chemical reactions - six catalysis, two degradations and one or two annihilation reactions.

### 1.1 Chemical reaction networks

We define a CRN as a set of reactions between chemical species *X*_*j*_. For example, in the reaction between species *X*_1_ and *X*_2_ represented by

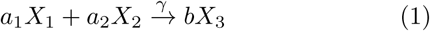

the reactants on the left are converted into the product *X*_3_ on the right at a rate *γ*, according to the stoichiometric coefficients *a*_1_, *a*_2_ and *b*. We model the evolution of the concentration *x*_*j*_ of the species *X*_*j*_ using mass action kinetics (Tóth and Érdi (1989)). For (1), the ordinary differential equations (ODEs) result in

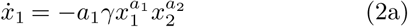

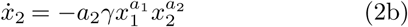

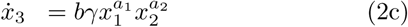

Mass action kinetics combined with dual rail representation enables the use of CRNs to model systems with both positive and negative signals. For example, the reactions

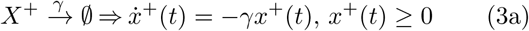

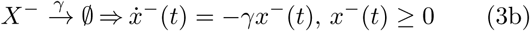

provide a representation of exponential decay of the real signal *x*(*t*) = *x*^+^(*t*) − *x*^−^(*t*), since

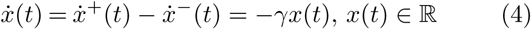

### 1.2 DNA strand displacement reactions

In a DSD reaction, the signal species are single stranded DNA molecules, composed of a binding domain and a toehold used to initiate strand hybridisation and tune the reaction rates. In Fig. 1, the hybridisation of the toehold 1 of the incoming strand to a complementary toehold 1* starts branch migration, displacing the domain 2, and releasing the output strands. The complete displacement is irreversible, due to the absence of overhanging complementary toeholds in the output strands. Given its irreversibility, and that the output strand can participate in other reactions, this enables a cascade of multiple reactions.

**Fig. 1.**
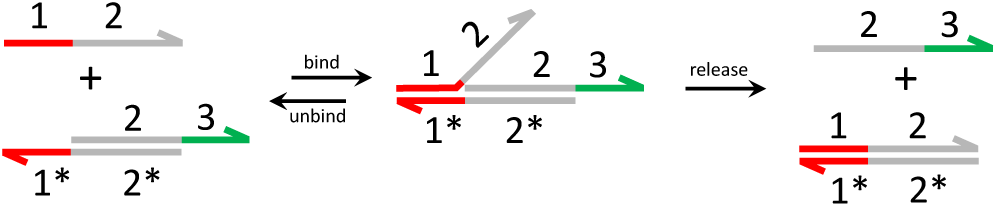
Illustration of a strand displacement reaction involving single and double strands, initiated by over-hanging toeholds.

As discussed in Oishi and Klavins (2011) and Chiu et al. (2015), with the dual rail representation it is possible to represent linear and feedback systems with CRNs composed of three types of reactions:

- catalysis 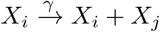
- degradation 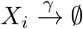
- annihilation 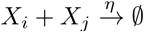

which have equivalent representations with DSD reactions (see for example Soloveichik et al. (2010), Chen et al. (2013), Srinivas et al. (2017), or Cherry and Qian (2018)).

For analysis and verification, we simulate the DSD circuitry by programming the species and affinities in Visual DSD (Lakin et al. (2017)), a specialised CAD tool for the simulation of strand displacement reactions. The simulations run in ‘default’ mode, where finite binding and unbinding rates are considered. The reaction rates in the CRNs are translated into affinities between the toeholds that mediate the initial hybridisation between the strands.

## 2. A MINIMALLY COMPLEX INTEGRAL FEEDBACK CONTROL SYSTEM

We now propose a representation of an integral feedback control system, using the fewer number of chemical reactions possible, followed by an implementation in Visual DSD using strand displacement reactions.

### 2.1 Integral control of a stable first order system

For this example we take the first order plant

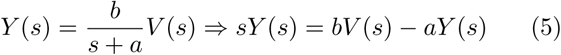

considering integral control action on the output error according to Fig. 2. The closed-loop dynamics are then

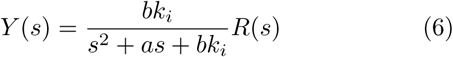

where the integral control ensures steady state tracking with *Y* (0) = *R*(0). The transient dynamics are defined by the roots of the characteristic polynomial 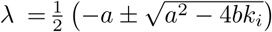, with a natural frequency 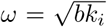 and damping coefficient 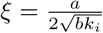.

**Fig. 2.**
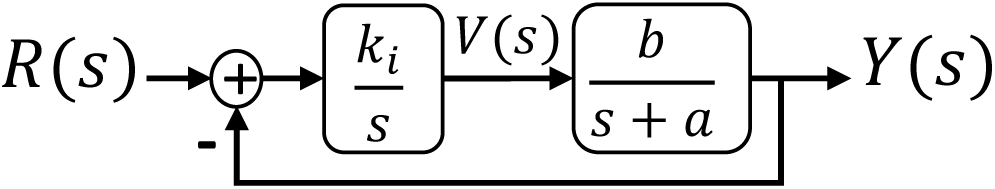
Reference tracking problem with integral control, of a stable first order plant.

### 2.2 Representation with chemical reactions

Fig. 3 shows a network of catalysis, degradation and annihilation reactions used to represent the closed-loop response, where

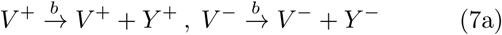

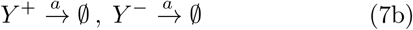

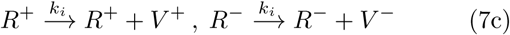

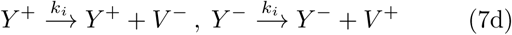

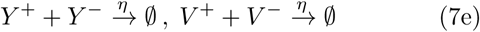

**Fig. 3.**
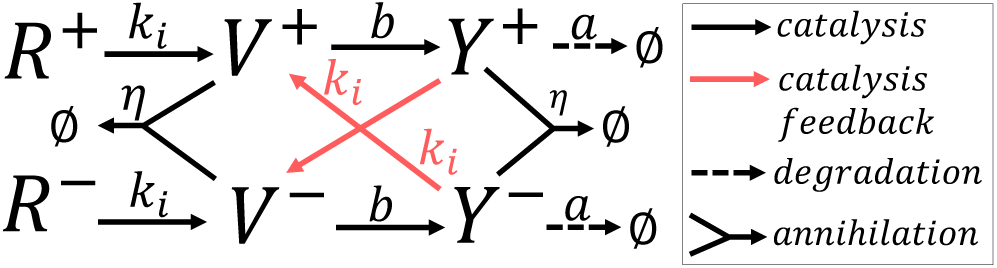
Representation of the integral control problem, with a chemical network of catalysis, degradation and annihilation reactions.

The reactions (7a-7b) represent the plant with gain *b* and a stable pole *s* = −*a*. Instead of using separate CRNs for subtraction and integration, both operations are combined to reduce the number of reactions. The additional dynamics used to compute the error in Paulino et al. (2019) are removed, and the contributions of the reference and output to the integral are subtracted in (7c-7d) by crossing the contributions from *Y* ^±^ to *V* ^∓^. The same reactions apply the control gain *k*_*i*_. The annihilation reactions in (7e) ensure low (i.e. experimentally feasible) concentrations of the chemical species, as discussed in Oishi and Klavins (2011).

Writing the respective mass action kinetics guiding the concentrations of the species, we have

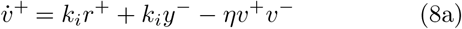

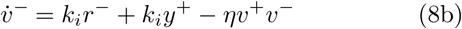

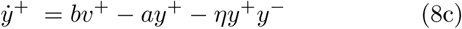

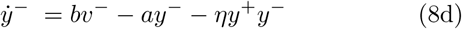

We have then for the dual rail quantities *y* = [*Y* ^+^] − [*Y* ^−^], *v* = [*V* ^+^] − [*V* ^−^], *r* = [*R*^+^] − [*R*^−^], that the input-to-output (I/O) dynamics are 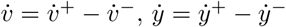, and

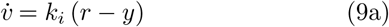

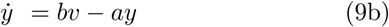

The I/O dynamics in (9) are linear and we recover the dynamics of the closed-loop system in (6). For simplification, any annihilation reaction between the reference species 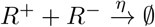 is disregarded, assuming the input concentrations are low. These will remain constant throughout the operation of the circuit, and the dynamics in (9) depend only on the difference *r* and not the concentration levels of *R*^+^ and *R*^−^.

### 2.3 Representation with strand displacement reactions

To translate the CRNs into nucleic acid chemistry we need sets of DSD reactions equivalent to each of the three types of reactions, and a mechanism to tune the reaction rates.

The catalysis and degradation reactions are set based on Join-Fork templates as in Chen et al. (2013). Fig. 4 shows examples of sets of DSD reactions equivalent to the unimolecular catalysis and degradation reactions introduced above, using auxiliary templates and intermediary strands.

**Fig. 4.**
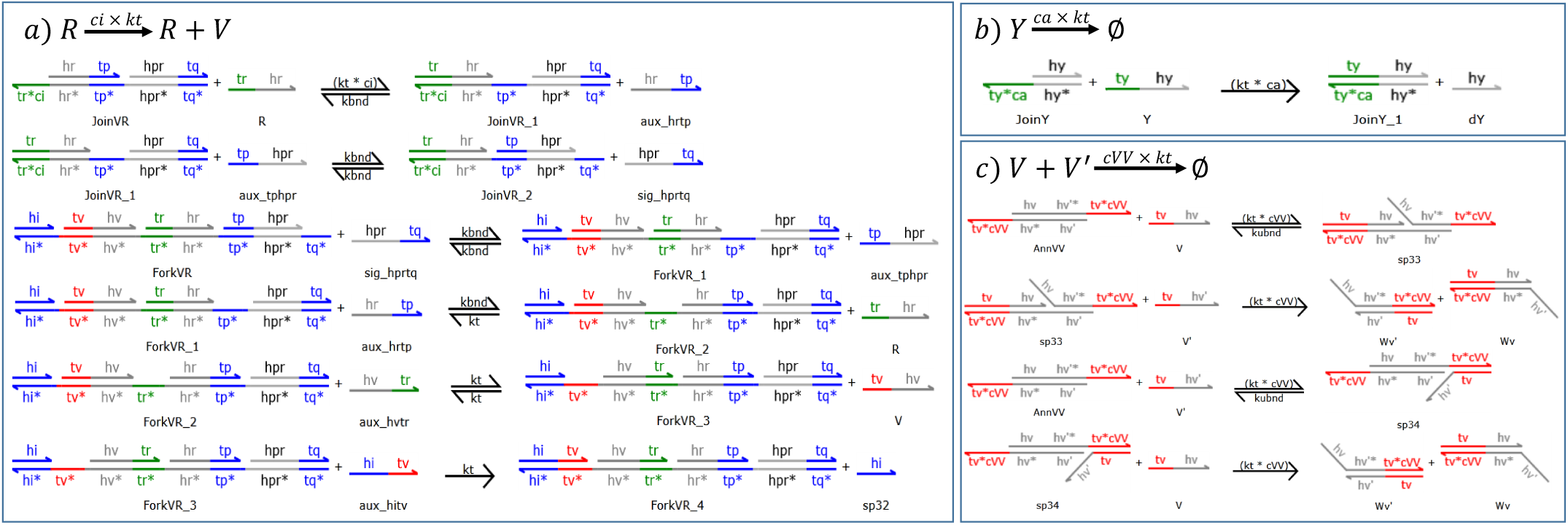
Examples generated in Visual DSD, of the DSD sets of reactions used to implement catalysis, degradation and annihilation.

Each signal species, such as *R* in Fig. 4a, is a single strand DNA containing toehold (<tr>) and binding (<hr>) domains. The toehold domain initiates hybridisation to the multi-stranded complex *JoinV R*, triggering a cascaded process to release an intermediary strand *sig*_*hprtq*. The availability of this new strand triggers a cascade of strand displacements involving the *ForkV R* complex, which releases the signal species *V* and returns a strand of *R* (according to the stoichiometry of *R* → *R* + *V*). The DSD reactions for the degradation in Fig. 4b are simpler, since the *JoinY* complex only needs to irreversibly capture the signal species *Y*. Since the bound complex does not have exposed toeholds, it becomes inert and no longer participates in the reactions.

The annihilation reactions are set with a single template per signal, following the cooperative hybridisation approach from Zhang (2011) and Cherry and Qian (2018). In the example shown in Fig. 4c, the irreversible capture of simultaneously present *V* and *V*′ is mediated by the presence of the template *AnnV V*. The presence of both *V* and *V*′ enables the second irreversible reaction into two waste double stranded complexes *Wv* and *Wv*′. The template complexes and auxiliary single stranded species are made available at a high concentration *C*_*max*_ to avoid their irreversible consumption from significantly impacting the dynamics.

The programmability of the DSD reactions results from the nucleotide sequences in the toeholds, which initiate strand hybridisation. As investigated in Zhang et al. (2018), the affinities between the base pairs define somewhat predictably the hybridisation kinetics, although, as described in Srinivas et al. (2017), other factors can also influence the effective reaction rate constants. For the purpose of our analysis, the rates of the DSD reactions are tuned by assigning degrees of complementarity between toeholds as in Yordanov et al. (2014) (see also Lakin et al. (2017)), where mismatches in the nucleotide sequences weaken the binding affinities. For example, the toehold <tr*ci> in the complex *JoinV R* in Fig. 4a has a degree of complementarity to the signal toehold <tr> of 0 *< ci* ≤ 1. If *kt* is the maximum binding rate between complementary toeholds <tr> and <tr*>, then the reaction mediated by <tr> and <tr*ci> is slowed down to *kt* × *ci*.

The implementation of the DSD reactions follows the six catalysis and two degradation from the CRN in (7), but with only one annihilation reaction. From the analysis with Visual DSD, we can remove the reaction *Y* ^+^ + *Y* ^−^→ ø to simplify further the implementation, since the concentrations of *Y* ^+^ and *Y* ^−^ remain sufficiently low.

The DSD reactions are set using the templates from Fig. 4, and the circuit is initialised with high concentrations of fifteen double stranded DNA templates and twenty single stranded auxiliary species. The simulation with Visual DSD is set with *C*_*max*_ = 10^5^ nM (auxiliary strands are consumed irreversibly and not replenished), a maximum toehold binding rate 10^−3^ (nMs)^−1^, and unbinding rate 0.1 s^−1^. See all the parameters in Table 1.

**Table 1.** Parameterisation for the Visual DSD simulation of the integral control circuit.

|  | Description | Values | Units |
| --- | --- | --- | --- |
| $C_{max}$ | Initial concentrations of template and auxiliary species | $10^5$ | nM |
| $kbnd$ | Toehold maximum binding rate for auxiliary strands ($\langle tp \rangle, \langle tq \rangle$) | $10^{-3}$ | (nMs)$^{-1}$ |
| $kubnd$ | Unbinding rate | 0.1 | (s)$^{-1}$ |
| $kt$ | Toehold maximum binding rate for signal species ($\langle tr \rangle, \langle ty \rangle, \langle tv \rangle$) | $10^{-4}$ | (nMs)$^{-1}$ |
| $ca$ | Toehold degree of complementarity for the degradation of $Y^\pm$ | $2.5 \times 10^{-3}$ | — |
| $cb$ | Toehold degree of complementarity for catalysis reaction in the plant | $10^{-3}$ | — |
| $ci$ | Toehold degree of complementarity for feedback catalysis implementing integral control | $5 \times 10^{-2}$ | — |
| $cVV$ | Toehold degree of complementarity for the annihilation reaction | 0.25 | — |

The simulations in Fig. 5 show that the reference tracking behaviour of the output *y* = [*Y* ^+^] − [*Y* ^−^] and evolution of the concentrations in the DSD reactions are in agreement with the output and concentrations of the CRN in (8) obtained by mass-action kinetics. The circuit based on DSD reactions shows the desired tracking behaviour, although the transient dynamics are more damped, probably due to the presence of the additional auxiliary species and bimolecular reactions.

**Fig. 5.**
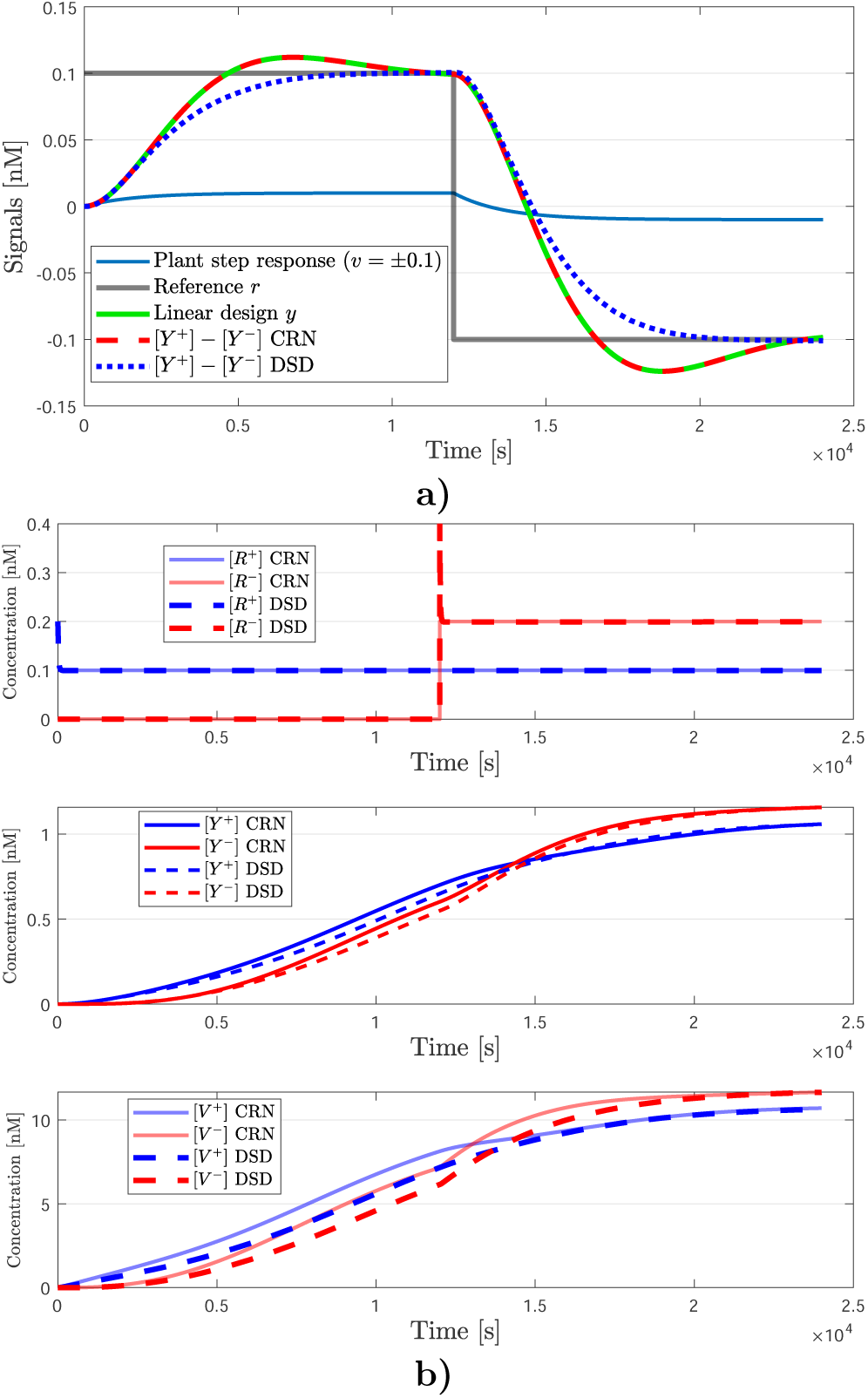
Simulations for the integral control example: a) comparison of the output *y* for the plant open-loop step response, closed-loop dynamics, CRN mass action kinetics and DSD reactions; b) comparison of the concentrations in the CRN and DSD representations.

## 3. A MINIMALLY COMPLEX STATE FEEDBACK CONTROL SYSTEM

In static state feedback, the controller modifies the closed-loop dynamics using only gains on the state of the plant and does not add dynamics to the open-loop system. For the plant, we use the simplest second order system, a double integrator. Besides output feedback, we thus have an extra state for feedback. The plant is more challenging than in the previous example, since it is marginally stable, with two poles at the origin, and thus besides the reference tracking requirement, the closed-loop system also needs to stabilise the open-loop system.

### 3.1 State feedback control of a double integrator

The process to be controlled is the classic double integrator

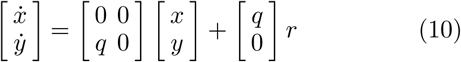

where each integration has a gain *q* (Fig. 6). With the negative state feedback control law *v* = *r* − *k*_1_*x* − *k*_2_*y*, we have two parameters to tune the dynamics of the closed-loop state space

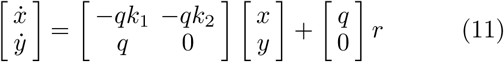

**Fig. 6.**
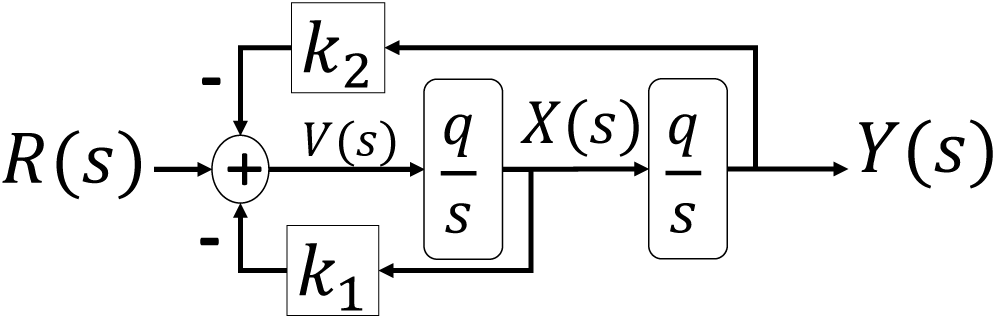
Block diagram representation of a plant (classic double integrator) stabilised by static state feedback.

The closed-loop frequency response results in a second order system with the transfer function

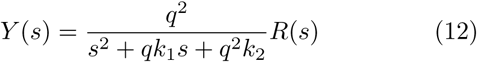

where the transient response is defined by the poles 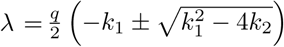.

The closed-loop system has only three parameters: one for the plant and two gains for the controller. From (12), we have *q*, which sets the timescale of the system, a static gain 1*/k*_2_, a natural frequency 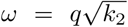, and damping coefficient 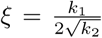. For steady state reference tracking, we need *k*_2_ = 1, which means that any error or deviation in this parameter introduced by implementation will impact the steady state error. It follows also that for an overdamped response ξ > 1 ⇒ *k*_1_ > 2.

### 3.2 Representation with chemical reactions

As in the previous example, the CRN representation is further simplified by combining the sum of the feedback contributions and reference with the integration of the first state. In this way, we avoid the additional dynamics of representing the sum with the steady state solution of additional reactions, as proposed in Paulino et al. (2019). Accounting for the dual rail representation, the CRN results in eight catalysis and two annihilation reactions, given by

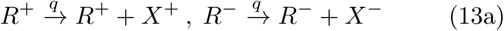

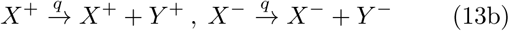

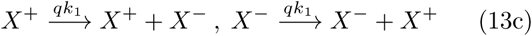

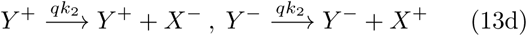

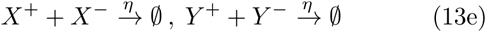

The chain of two catalysis reactions (13a-13b) represent the double integrator in the plant. The marginal stability of the integration can be related with the marginal stability of the stoichiometry in the catalysis reactions, where the produced species need to be bound for stability. The reactions in (13c-13d) implement the negative gains, by crossing the contributions between the dual rail species (Fig. 7a). Finally, the annihilation reactions are put in place in (13e), to ensure the concentrations are kept at feasible levels.

**Fig. 7.**
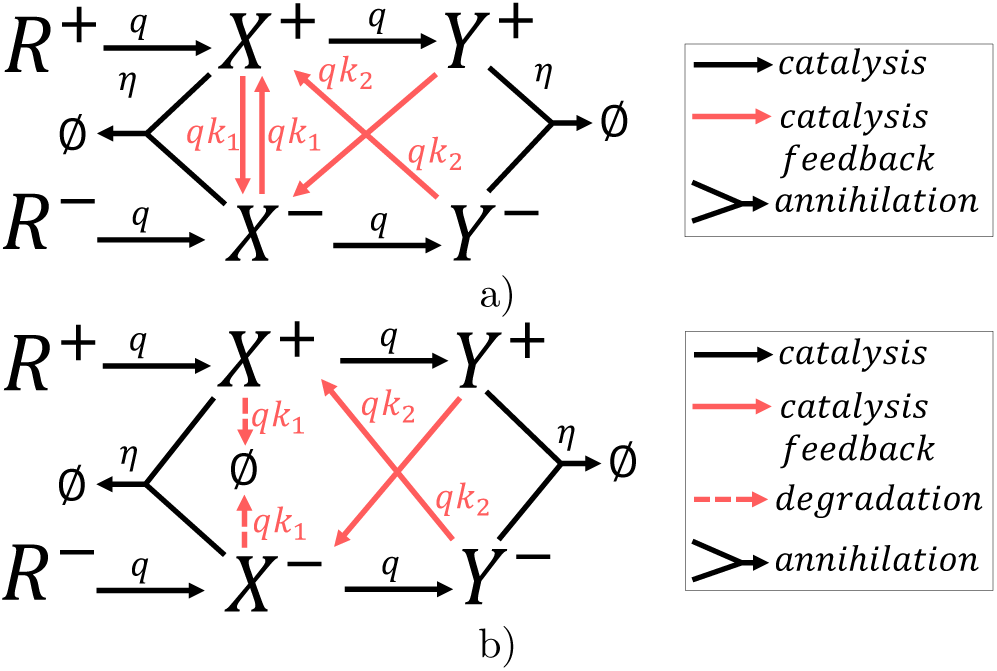
Network of chemical reactions with dual rail representation, using a) *catalytic degradation* or b) degradation reactions.

The mass action kinetics for the chemical network results in

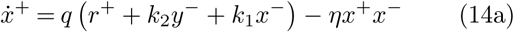

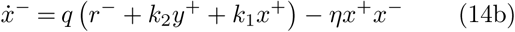

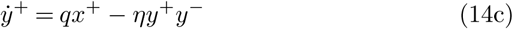

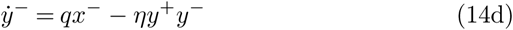

From the reversed contributions in (14a-14b), the negative gains in the I/O dynamics of 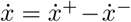 and 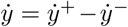 are given by

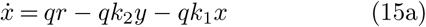

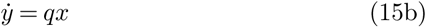

In (15) we recover the linear closed-loop dynamics.

The combined effect of crossed catalysis reactions 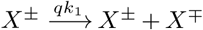 and the fast annihilation 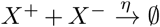 results in the *catalytic degradation* proposed in Yordanov et al. (2014). Hence, the autorepressing gain is the same as 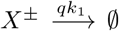 (for fast *η* ≫ *qk*_1_), and alternatively we can replace the catalysis in (13c) with degradation reactions. This results in the CRN from Fig. 7b with six catalysis, two degradation, and two annihilation reactions given by

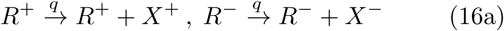

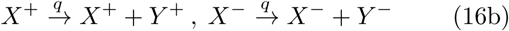

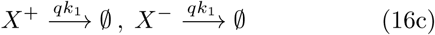

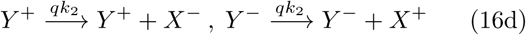

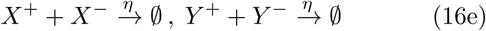

The mass action kinetics are now different, with

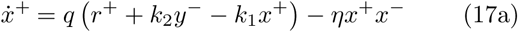

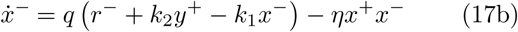

but the I/O dynamics of 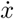 are the same as in (15a). The CRN representation is compared with the linear design in Fig. 8, showing the prescribed reference tracking behaviour, and the matching between the linear control design and the trajectories of the dual rail signals resulting from I/O dynamics of the CRN.

**Fig. 8.**
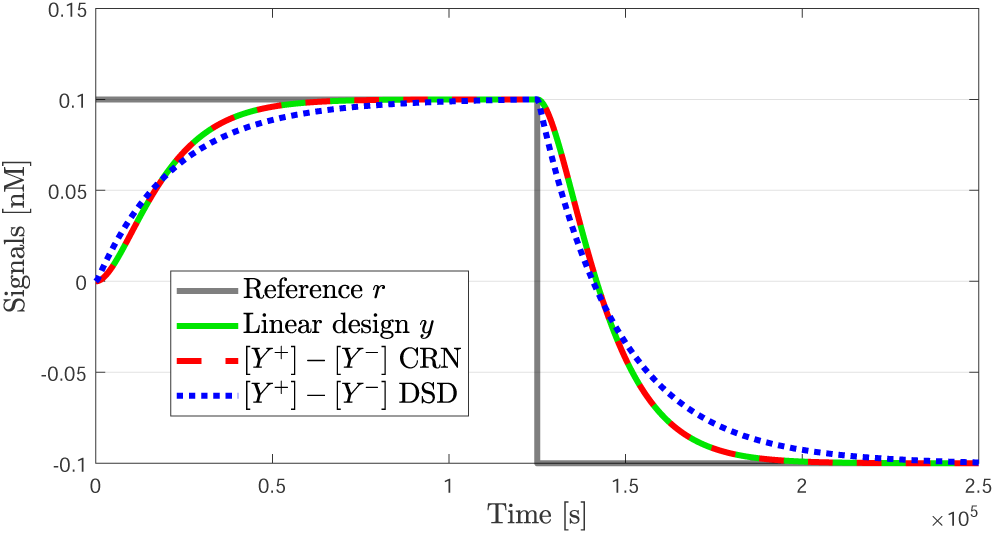
Steady state tracking response of the linear design and the I/O dynamics of the representations with chemical reactions and DSD reactions.

### 3.3 Representation with strand displacement reactions

For the DSD representation, the crossed feedback between *X*^+^ and *X*^−^ is replaced by degradation reactions, and the architecture from Fig. 4 is applied to obtain DSD reactions equivalent to the CRN in (16). Furthermore, from the analysis with Visual DSD, the circuit can operate without the annihilation reaction 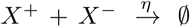, since these concentrations remain low. Depending on the experimental set-up, this is yet another possible simplification.

With these simplifications, although we have state feedback of a more complex marginally stable second order system and more degrees of freedom, the implementation has the same level of complexity as the previous integral control problem, with six catalysis, two degradation, and one annihilation reactions. The simulation in Visual DSD has fifteen double stranded complexes and twenty auxiliary single stranded species initialised at *C*_*max*_ = 10^5^ nM (irreversibly consumed without replenishment), with a maximum toehold binding rate 10^−3^ (nMs)^−1^, and unbinding rate 0.1 s^−1^. See all the parameters in Table 2.

**Table 2.** Parameterisation for the Visual DSD simulation of the state feedback circuit.

|  | Description | Vales | Units |
| --- | --- | --- | --- |
| $C_{max}$ | Initial concentrations of template and auxiliary species | $10^5$ | nM |
| $k_{bnd}$ | Toehold maximum binding rate for auxiliary strands (<tp>,<tq>) | $10^{-3}$ | (nMs)$^{-1}$ |
| $k_{ubnd}$ | Unbinding rate | 0.1 | (s)$^{-1}$ |
| $kt$ | Toehold maximum binding rate for signal species (<br>,<ty>,<tx>) | $10^{-4}$ | (nMs)$^{-1}$ |
| $cq$ | Toehold degree of complementarity for reactions in the plant | $8 \times 10^{-3}$ | — |
| $c1$ | Toehold degree of complementarity for the degradation of $X^\pm$ | $8 \times 10^{-3}$ | — |
| $c2$ | Toehold degree of complementarity for feedback catalysis of $Y^\pm$ | $8 \times 10^{-3}$ | — |
| $cYY$ | Toehold degree of complementarity for the annihilation reaction | 1 | — |

The DSD circuit shows reference tracking behaviour (Fig. 8) while the I/O dynamics of the CRN matches well the linear design, the transient dynamics with the DSD reactions are slower. Comparing the CRN concentrations with the concentrations in the Visual DSD simulation, we also see slower dynamics for *Y* ^±^ and lower equilibrium levels for *X*^±^, indicating a higher damping of the state *x* (Fig. 9). Although further tuning is necessary, the dynamics are close to the desired state feedback design, and also indicate that indeed the annihilation reaction for *X*^±^ may not be necessary due to the limiting degradation.

**Fig. 9.**
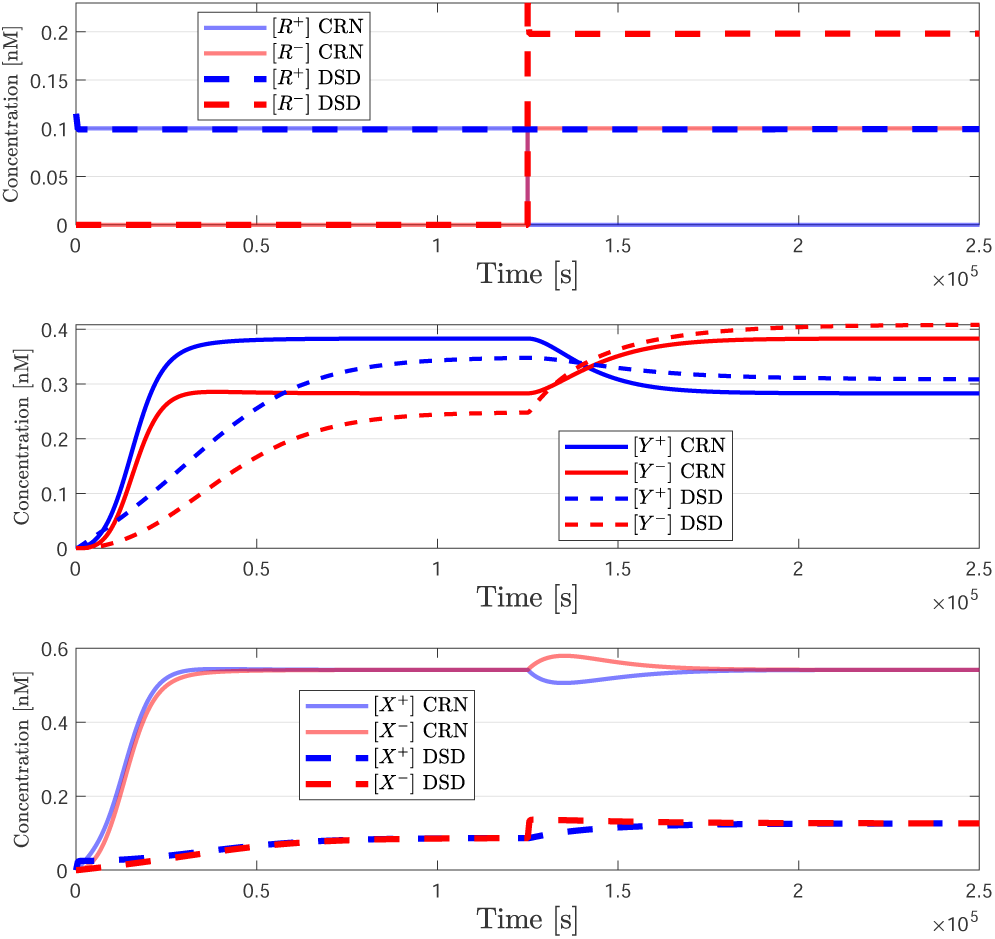
Comparison of the concentrations for the abstract CRN, and the DSD implementation. The steady state conditions for the output species *Y* ^±^ are similar in the CRN and DSD representations, although smaller for *X*^±^ in the DSD circuit.

For the DSD network, we adopted the construction in (16), where catalysis reactions responsible for the state feedback are replaced with the degradation reactions in (16c), which use the simpler template complexes in Fig. 4b, and fewer species. However, there may be advantages in building the circuit relying only on catalysis reactions (and annihilation). When introducing the catalytic degradation scheme, Yordanov et al. (2014) argue that spatial localisation of the catalysis reactions could allow faster degradation rates.

## 4. CONCLUSIONS

There currently exist mature theoretical frameworks for the design of linear feedback controllers with chemical reaction networks. Systematic procedures and software tools provide a translation to equivalent reactions based on nucleic acid chemistry, which should be amenable to experimental implementation. However, the readiness level of this technology has not followed the theoretical developments, and we are lacking experimental validation of such systems.

We propose here two control problems, which can be represented with very few CRN and DSD reactions, as minimally complex candidates for future experimental validation of feedback circuits using strand displacement reaction networks. The reduced number of reactions puts these DSD networks within the current capabilities for experimental investigation, while still capturing important features of general linear feedback control systems.

Although simple, the examples are representative of standard control designs, and the circuits are interesting for future experimental investigation of the dependence of closed-loop dynamics on toehold design, and integration of the annihilation reactions. Future experimental testing could also give insight into the effects of crosstalk and leakage on feedback stability and performance, and provide data for developing sharper uncertainty models for taking these effects into account during the design process.

## REFERENCES

Buisman, H.J., ten Eikelder, H.M.M., Hilbers, P.A.J., and Liekens, A.M.L. (2008). Computing algebraic functions with biochemical reaction networks. Artificial Life, 15(1), 5–19.

Chatterjee, G., Dalchau, N., Muscat, R.A., Phillips, A., and Seelig, G. (2017). A spatially localized architecture for fast and modular DNA computing. Nature Nanotechnology, 12(9), 920–927.

Chen, Y.J., Dalchau, N., Srinivas, N., Phillips, A., Cardelli, L., Soloveichik, D., and Seelig, G. (2013). Programmable chemical controllers made from DNA. Nature Nanotechnology, 8(10), 755–762.

Cherry, K.M. and Qian, L. (2018). Scaling up molecular pattern recognition with DNA-based winner-take-all neural networks. Nature, 559(7714), 370–376.

Chiu, T.Y., Chiang, H.J.K., Huang, R.Y., Jiang, J.H.R., and Fages, F. (2015). Synthesizing configurable biochemical implementation of linear systems from their transfer function specifications. PLoS ONE, 10(9), e0137442.

Dannenberg, F., Kwiatkowska, M., Thachuk, C., and Turberfield, A.J. (2015). DNA walker circuits: computational potential, design, and verification. Natural Computing, 14(2), 195–211.

Foo, M., Kim, J., Sawlekar, R., and Bates, D.G. (2017). Design of an embedded inverse-feedforward biomolecular tracking controller for enzymatic reaction processes. Computers & Chemical Engineering, 99, 145–157.

Hemphill, J. and Deiters, A. (2013). DNA computation in mammalian cells: microRNA logic operations. Journal of the American Chemical Society, 135(28), 10512–10518.

Joesaar, A., Yang, S., Bögels, B., van der Linden, A., Pieters, P., Kumar, B.V., Dalchau, N., Phillips, A., Mann, S., and de Greef, T.F.A. (2019). DNA-based communication in populations of synthetic protocells. Nature Nanotechnology, 14(4), 369–378.

Lakin, M.R., Petersen, R.L., and Phillips, A. (2017). Visual DSD user manual v0.14 beta.

Oishi, K. and Klavins, E. (2011). Biomolecular implementation of linear I/O systems. IET Systems Biology, 5(4), 252–260.

Paulino, N.M.G., Foo, M., Kim, J., and Bates, D.G. (2019). PID and state feedback controllers using DNA strand displacement reactions. IEEE Control Systems Letters, 3(4), 805–810.

Qian, L. and Winfree, E. (2011). Scaling up digital circuit computation with DNA strand displacement cascades. Science, 332(6034), 1196–1201.

Seelig, G., Soloveichik, D., Zhang, D.Y., and Winfree, E. (2006). Enzyme-free nucleic acid logic circuits. Science, 314(5805), 1585–1588.

Soloveichik, D., Seelig, G., and Winfree, E. (2010). DNA as a universal substrate for chemical kinetics. Proceedings of National Academy of Sciences, 107(12), 5393–5398.

Srinivas, N., Parkin, J., Seelig, G., Winfree, E., and Soloveichik, D. (2017). Enzyme-free nucleic acid dynamical systems. Science, 358(6369), eaal2052.

Thubagere, A.J., Thachuk, C., Berleant, J., Johnson, R.F., Ardelean, D.A., Cherry, K.M., and Qian, L. (2017). Compiler-aided systematic construction of large-scale DNA strand displacement circuits using unpurified components. Nature Communications, 8, 14373.

Tóth, P. and Érdi, J. (1989). Mathematical models of chemical reactions: Theory and Applications of Deterministic and Stochastic Models. Manchester University Press.

Wang, B., Thachuk, C., Ellington, A.D., and Soloveichik, D. (2017). The design space of strand displacement cascades with toehold-size clamps. In International Conference on DNA-Based Computers, 64–81. Springer.

Wang, B., Thachuk, C., Ellington, A.D., Winfree, E., and Soloveichik, D. (2018). Effective design principles for leakless strand displacement systems. Proceedings of the National Academy of Sciences, 115(52), E12182–E12191.

Yordanov, B., Kim, J., Petersen, R.L., Shudy, A., Kulkarni, V.V., and Phillips, A. (2014). Computational design of nucleic acid feedback control circuits. ACS Synthetic Biology, 3(8), 600–616.

Zhang, D.Y. (2011). Cooperative hybridization of oligonucleotides. Journal of the American Chemical Society, 133(4), 1077–1086.

Zhang, J.X., Fang, J.Z., Duan, W., Wu, L.R., Zhang, A.W., Dalchau, N., Yordanov, B., Petersen, R., Phillips, A., and Zhang, D.Y. (2018). Predicting DNA hybridization kinetics from sequence. Nature Chemistry, 10(1), 91–98.

